# Testing Hormone-specific Antibody Probes for Presumptive Detection and Separation of Contributor Cell Populations in Trace DNA Mixtures

**DOI:** 10.1101/2019.12.18.881748

**Authors:** Jennifer M. Miller, Emily R. Brocato, Vamsi K. Yadavalli, Susan A. Greenspoon, Christopher J. Ehrhardt

## Abstract

“Touch” or trace DNA evidence represent a significant proportion of samples analyzed by forensic science laboratories. Because these samples frequently contain multiple contributors and are often challenging to analyze due to low DNA concentrations and frequent degradation, front end techniques to simplify the mixture prior to DNA profiling could significantly impact case processing and enhance success rates. The goal of this study was to investigate whether targeting hormone molecules within the cell with antibody probes could be used to selectively label and then physically isolate contributor cell populations in trace biological samples. The separation of male and female cells into distinct fractions could reduce the complexity of the mixture prior to DNA profiling. To accomplish this, we first tested the specificity of fluorescently labelled anti-testosterone and anti-dihydrotestosterone antibody probes to epidermal cells from both male and female individuals. Results show that male and female cell populations can be effectively labeled using anti-testosterone and anti-dihydrotestosterone antibody probes and that distinct differences in binding efficiency and resulting median fluorescence of cell populations were observed between several individuals. These differences were then used to design sorting criteria for physically isolating each cell population in two-person epidermal cell mixtures using fluorescence activated cell sorting (FACS). DNA profiling of separated fractions in combination with probabilistic modeling demonstrated that some cell mixtures could be enriched for one contributor in separated cell fractions and yielded statistically more discriminating profiles compared to those generated from the original mixtures. Other mixtures tested showed less evidence of effective cell separation possibly due to a number of factors including imbalance of contributor DNA ratio, intra-sex variation of antibody binding efficiency, and contributions of extracellular or cell-free DNA in the mixture sample. Screening and separation of trace DNA samples with this approach may be presumptive and ultimately constrained by specific parameters of the original mixture, however, antibody binding optimization may mitigate some of these influences.

## Introduction

There is a strong interest in the development of strategies for separating cell populations from mixtures *prior* to DNA profiling. The earliest systems focused on sperm and epithelial cell mixtures beginning with the advent of the differential extraction procedure (1,2). Recently, techniques have been described for whole blood mixtures (3,4), and blood/buccal mixtures (5). Although mixtures comprised solely of shed epidermal cell populations (i.e., touch/trace cell mixtures) are a priority for forensic investigators, relatively few approaches have been demonstrated for this sample type (6). This is likely due to the unique characteristics of touch epidermal cell populations compared to other sources of tissues. Cells deposited through contact are derived from keratinized, stratified squamous epithelial tissue (7) and specifically, the outer layer of the epidermis, known as the stratum corneum. This layer can have as many as 30 layers of dead cells (8) and may approach ~40μm in thickness. As cells are shed, they are replaced by younger keratinocytes that have undergone differentiation as they migrate from the deepest layer (stratum basale) through the epidermis (7,9).

An important implication of the differentiation process is that prior to shedding, epidermal cells have lost a significant portion of their intracellular contents including the nucleus and other organelles, genomic DNA, and various cell-specific or tissue specific antigen groups (10). Thus, many of the traditional molecular targets used to differentiate cell populations in a mixture are either absent or inaccessible in trace biological samples. In order to develop a front-end separation method for touch/trace mixtures, new molecular targets are therefore needed. Hormone molecules that are differentially expressed across sexes are one such promising system. For instance, intracellular levels of testosterone and dihydrotestosterone (DHT) in serum are 10-30x greater in adult males compared to females (11–14). Although there is little data on the abundance of hormone molecules specifically within epidermal cells, testosterone and dihydrotestosterone are pervasive signaling molecules in different tissues, including epithelial cells, throughout the body (13–15). This general disparity in hormone concentration between male and female epithelial cells make molecules such as testosterone promising targets for antibody-based cell tagging approaches, to label and then isolate male contributor cell populations within a mixture prior to DNA analysis. Additionally, testosterone levels of stored serum have been found to be consistent for over 40 years, supporting its potential as a stable biomarker for forensic samples, which may frequently be compromised by varying degrees of environmental exposure prior to collection (16).

In this study, we test the binding efficiency of antibody probes targeting hormone molecules for labeling cell populations composed of only shed epidermal cells from male and female individuals. We examine both the consistency of antibody interactions in labelling cells from different individuals and also assess whether different patterns of antibody interaction are evident between male and female contributors. Fluorescently labeled antibody probes specific to testosterone and DHT were used. Labeled epithelial skin cell populations were then analyzed using flow cytometry to characterize the efficiency of antibody probe binding to the respective cell populations. Finally, we test the potential utility of hormone-specific antibody probes for separating contributor fractions of trace DNA mixtures by physically separating labeled mixtures using Fluorescence Activated Cell Sorting (FACS) followed by STR typing of each separated fraction. Quantitative assessment of the degree of enrichment was performed for the male and female contributors in the post-sort fractions by use of probabilistic modeling.

## Materials and Methods

### Sample Collection

Epithelial skin cells were collected from male and female volunteers using a Whatman^®^ FTA® Sterile Omni Swab (GE Healthcare, Chicago, IL). Participants were asked to swab the sides of their nose and behind their ears for approximately thirty seconds using swabs in order to maximize skin cell yield. Participants who were wearing makeup, or any other facial-care product, were asked to only swab behind their ears. Informed consent was obtained from the participants prior to performing the experiments.

Swabs from each individual were then placed into a single tube containing 2mL of cell staining buffer (BioLegend, San Diego, CA). To elute the cells from the swabs, the swabs were incubated in buffer for approximately five minutes and then pulse agitated in a vortex platform (300rpm, 10 seconds).

### Antibody Staining and Flow Cytometry Screening

For antibody staining experiments, two swabs were collected from each participant. Both swabs were placed into a single tube containing 2 mL of cell staining buffer to elute cells into a single solution. The cell solution was then centrifuged at 10,000xg for 10 minutes. The supernatant was then decanted so that 100 μL of buffer remained. Next, 1 μL of blocking buffer was added to the cell suspension (Thermo Fisher Scientific, Waltham, MA). The resulting cell suspension was incubated for 10 minutes on ice. After incubation, 2.5 μL of FITC-conjugated anti-testosterone antibody (Novus Biologicals, Littleton, CO) and 2.5 μL of FITC-conjugated anti-dihydrotestosterone antibody (Biomatik, Wilmington, DE) were added to the cell solution. This was then mixed and incubated on ice for one hour. After incubation, the cell solution was washed twice with cell staining buffer. A 50 μl aliquot of the initial cell solution was removed prior to antibody hybridization and was analyzed without further treatment as a negative control cell population for each experiment. Initial evaluations of hybridization and optimizations were performed on each antibody separately using either flow cytometry or fluorescence microscopy.

Initial screening and analysis of antibody hybridization efficiency on cell populations was performed using the Guava^®^ easyCyte^™^ flow cytometer (Millipore Inc., Burlington, MA). This information was used to assess antibody binding efficiency and the degree of separation between male and female cell populations. Epithelial cells in cell staining buffer were analyzed in the Guava^®^ easyCyte^™^ instrument in aliquots of 500 μL, after being passed through a 12×75 mm polystyrene filter. The FITC-labeled antibodies were excited with a 488 nm laser (50 mW).

### Imaging Flow Cytometry (IFC)

A subset of sample cell populations were analyzed using an Amnis^®^ Imagestream X Mark II (EMD Millipore; Burlington, MA) equipped with 405nm, 488nm, 561nm, and 642nm lasers. Laser voltages for tests were set at 120mW, 100mW, 100mW and 150mW, respectively. Images of individual events were captured in five detector channels labeled: 1 (430-505nm), 2 (505-560nm), 3 (560-595nm), 5 (640-745nm), and 6 (745-780nm). Channel 4 was used to capture images under brightfield illumination. Magnification was set at 40x and autofocus was enabled so that the focus varied with cell size. Raw image files (.rif) were then imported into IDEAS^®^ Software (EMD Millipore; Burlington, MA) to generate image galleries. The scale of fluorescence intensities displayed was set to the range of pixel values for each channel.

### Fluorescence Activated Cell Sorting (FACS)

Cell separation was performed on the Aria-BD FACSAria^™^ II High-Speed Cell Sorter (Beckton Dickinson, Franklin Lakes, NJ) using the 488 nm excitation laser. The laser voltages were set to 203 V for Side Scatter, and 475 V for 488 530/30 (FITC). Prior to flow cytometry analysis, the cell mixtures were passed through a 12×75 mm polystyrene filter (Falcon^®^, Corning, NY) to remove debris. Antibody hybridization and cell sorting was performed on 1:1 (vol:vol) mixtures of male and female epidermal cells (500μl each). Data obtained from the FACS instrument was analyzed using FACSDiva v. 6.1.3 software program (Becton Dickinson). For two mixtures, a small aliquot (50μl) of each individual donor cell solution was retained and analyzed separately on the cell sorter to assess hybridization efficiency as well as relative cell counts of the donor cell population prior to creating cell mixtures. For two additional mixtures (‘blind’/unknown mixture samples), analysis of donor cell populations was not performed prior to sorting of the mixture.

Sorting was performed for approximately one hour on each mixture sample so that a minimum of 400 cell ‘events’ could be collected per cell fraction. The amount of time it took to collect the requisite number of cells and conversely the amount of volume consumed during sorting did vary between mixture samples likely owing to differences in the concentration of cells and/or biological/non-biological debris. However, for all the mixture samples tested, less than ~50 μl of the original cell mixture remained following sorting. In order to assess the proportion of the original cell population that was captured into any sorted cell fraction and subsequently processed for DNA profiling, we compared the total number of cells collected across all fractions (i.e., right, middle, left) to the total number of cells detected with optical properties consistent with shed epidermal cells. For the four mixtures tested, the proportion of sorted cells ranged between ~50% and ~85% of the original cell population. Although it was not an explicit goal of this study, we note that any cells and/or biological material not sorted could nonetheless be collected and used for other analyses.

### STR Profiling of FACS Fractions

All sorted cell fractions as well as unsorted mixture samples were extracted manually with the DNA IQ^™^ System (Promega, Madison, WI) following the Virginia Department of Forensic Science’s protocol (17). Purified DNA extracts were concentrated from approximately 35 μL to ~13 μL using vacuum centrifugation (SpeedVac, Thermo Fisher Scientific). Quantitation was performed on all donor reference samples purified with Promega’s Plexor ^®^

HY System on a MX3005P^™^ Quantitative PCR instrument (Stratagene, Santa Clara, CA), which was equipped with Plexor Analysis software. Donor reference samples were comprised of either buccal swabs or epithelial skin swabs (from the forearm) from each individual. Additionally, for the first two mixture samples, DNA extracts were quantified using the same protocol as the reference samples. However, due to low DNA yields (<200pg), the DNA extracts for the following pre-sort and post sort samples was used at maximal volume (10 μL per VDFS protocol) for subsequent amplification steps without quantitation.

STR amplification was performed on the GeneAmp 9700 thermal cycler (Applied Biosystems ((ABI), Carlsbad, CA), using Promega’s PowerPlex^®^ Fusion System following the manufacturer’s recommendations. Ten microliters of DNA extract was added to 15 μL of the STR reaction mix for a full volume, 25 μL amplification reaction. STR products were separated on a 3500xl Genetic Analyzer (ABI) with a 24-second injection, followed by STR analysis with the GeneMapper ^®^ ID-X v1.4 software program (ABI), following the manufacturer’s recommended procedure. The analytical threshold used to interpret the STR profiles manually were 75 RFU for each dye channel. Probabilistic genotype modeling analysis was conducted using the TrueAllele^®^ Casework system (Cybergenetics, Pittsburgh, PA). The procedure was performed as described in the TrueAllele^®^ Casework (TA) user manuals and in (18,19). The qualitative, assessment of STR profiles was performed following the Virginia Department of Forensic Science’s protocols (17), however alleles were noted that were below the analytical threshold, but clearly distinguishable from baseline noise since they would be modeled by the TA system. Electropherograms for all DNA profiles are included as supplemental data with this manuscript.

## Results

### Binding efficiency of testosterone antibody probes in epidermal cells

Initial screening of epidermal cells for antibody probe binding was conducted on samples from 10 males and 10 females. Fluorescence histograms of the cell populations are shown in Figures 1 and 2. Probe binding was observed across all samples as evidenced by shifts in median fluorescence between the stained and unstained cell populations. The distribution of fluorescence values for each cell population showed significant variation between individuals with no clear systematic differences between male and female individuals (e.g., comparing stained histograms across Figures 1 and 2). However, several distinct subpopulations of cells with higher median fluorescence and minimal overlap with other fluorescence histograms were observed (e.g., subpopulations highlighted with asterisk in Figure 1).

**Figure 1.**
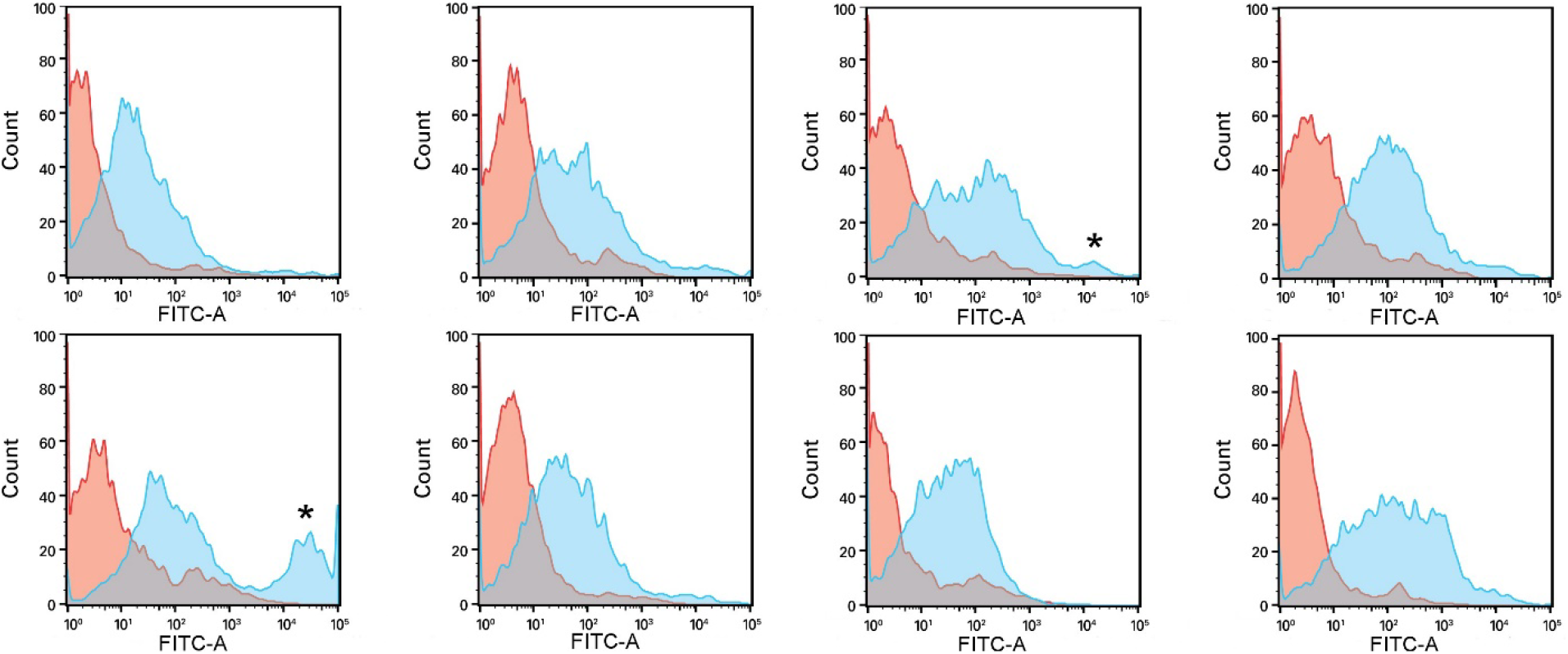
Fluorescence histograms of epithelial skin cell populations hybridized to anti-testosterone and anti-dihydrotestosterone antibodies (blue) and unstained (red) from the same contributor. Histograms from first two columns represent cell populations sampled from four male contributors and histograms in the right two columns were sampled from four female contributors. The X-axis shows fluorescence intensity, while the Y-axis shows the number of events analyzed for each sample. Fluorescence histograms from all contributors used for this study (10 males, 10 females) are provided in Figures S1, S2.

**Figure 2.**
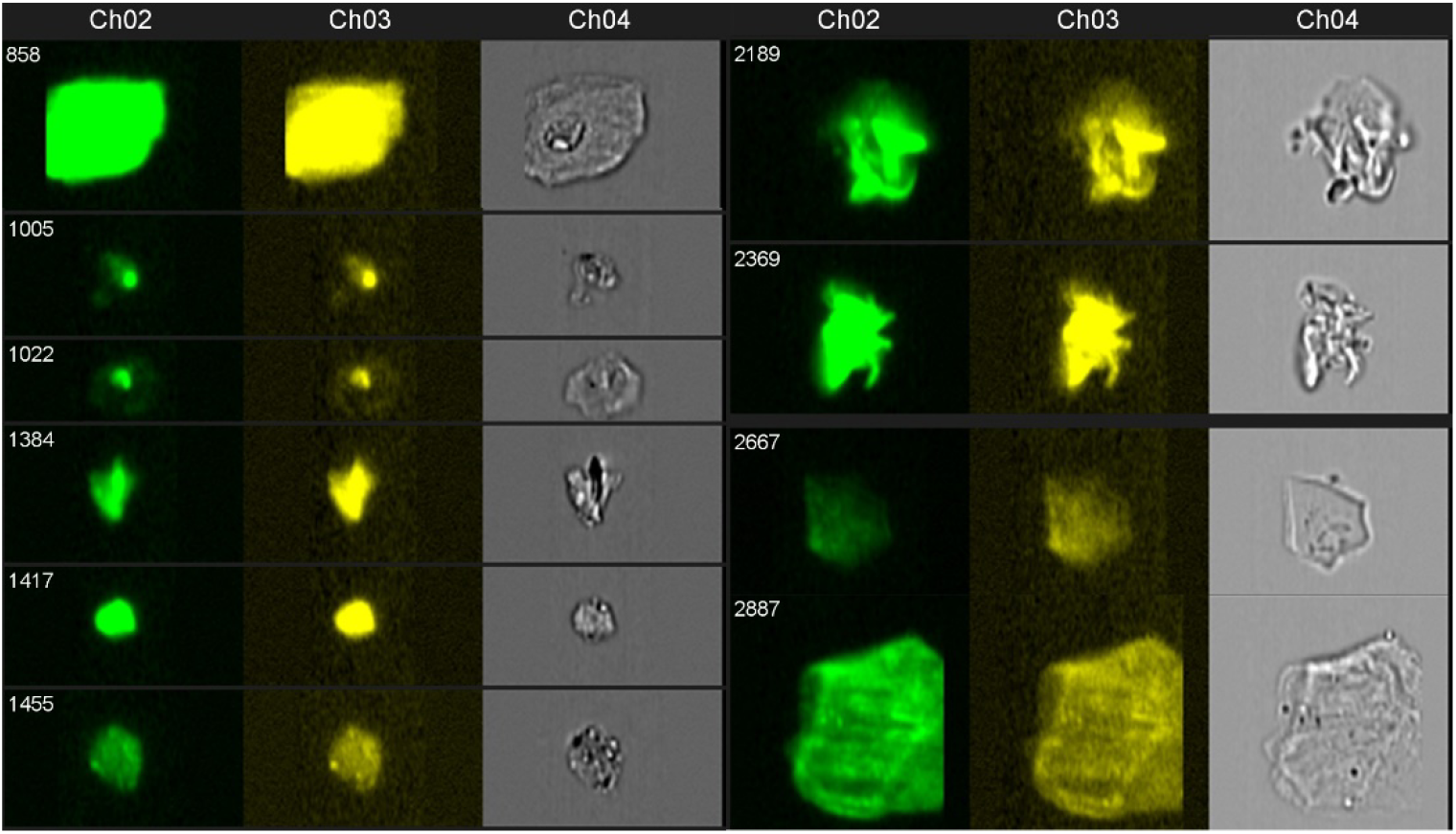
Image galleries of events analyzed from male (left) and female (right) epidermal cell populations after labeling with testosterone-specific antibody probe.

Antibody hybridization in epidermal cell samples was also analyzed using imaging flow cytometry. Figure 2 shows both brightfield and fluorescence imaging of individual epidermal cells that have been hybridized with anti-testosterone and anti-dihydrotestosterone probes: male (left) and female (right). For both male and female cell populations, fluorescence is observed with cell events that have a size and morphology consistent with epidermal tissue. However, the heterogeneity of cell sizes and morphologies indicate that cell debris, fragments, or other biological material may also be included in these cell populations (Figure 2, S3).

### Fluorescence Activated Cell Sorting and DNA Profiling: Mixture 1

Although antibody binding and resulting fluorescence profiles did not strictly correlate with the sex, it is important to note that the results indicate the possibility to differentiate contributor cell populations in some epidermal cell mixtures based on the efficacy of hybridization to testosterone-and DHT-specific probes. To determine how such differences can be used to facilitate a mixture separation workflow, equal volume mixtures of male to female cells were formed and hybridized to testosterone-and dihydrotestosterone-specific antibody probes. The contributor cell ratio was assessed by analyzing each donor cell solution separately prior to processing the mixture sample. For Mixture 1, the cell ratio was ~1.5: 1 (M:F). The epidermal cells in the two mixtures were incubated with anti-testosterone and anti-dihydrotestosterone antibodies and physically sorted into one of three fractions: ‘left’, ‘middle’, and ‘right’ (Figure 3, left panel).

**Figure 3.**
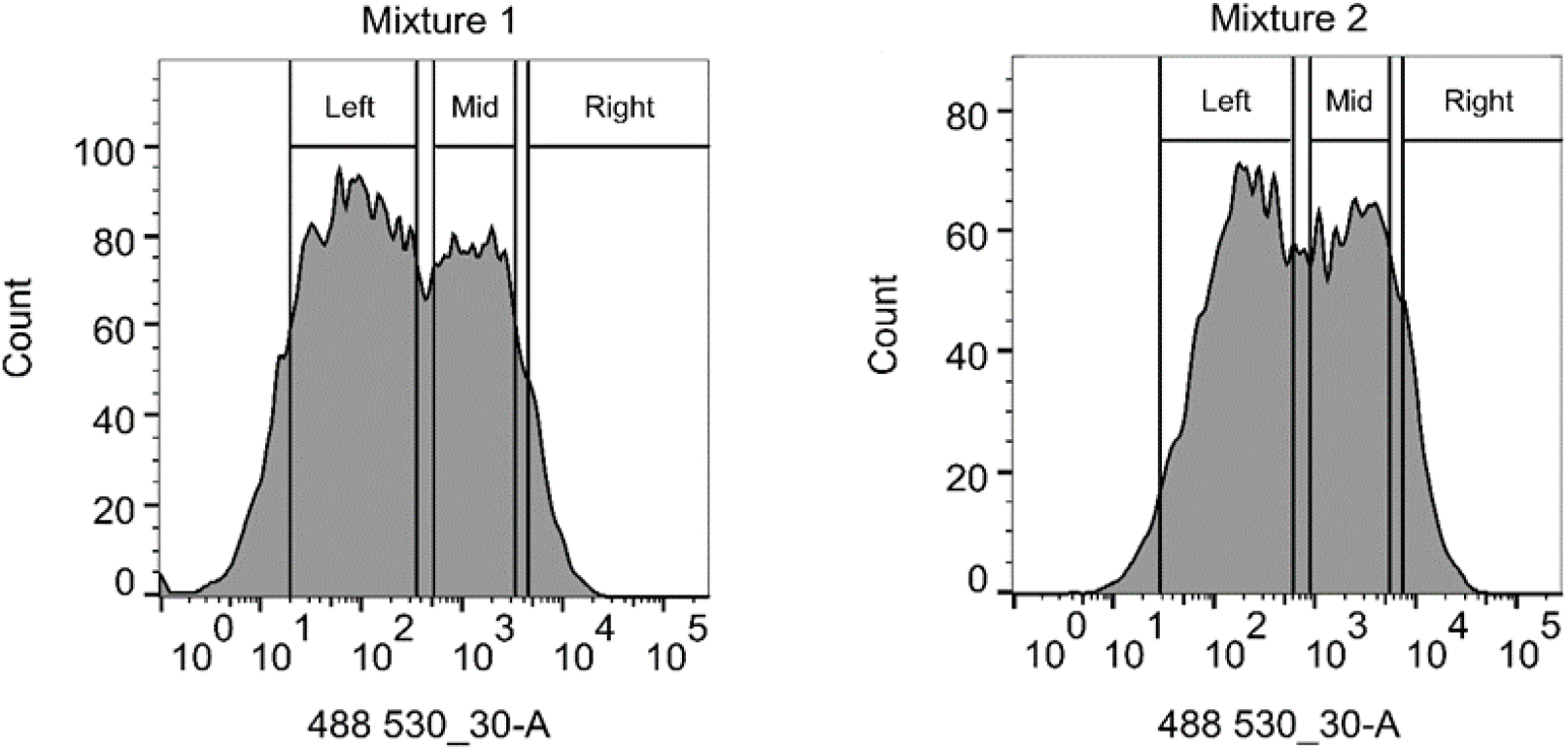
Gating parameters for fluorescence activated cell sorting on the Aria-BD FACSAria^™^ II High-Speed Cell Sorter for Mixtures 1 and 2.

The gating criteria for the first cell mixture was determined by an initial screening of the male and female donor reference profiles after antibody probe hybridization. As shown in earlier works, the position of the sorting gate is likely to have a significant effect on the number of cells collected for a given period of time, and conversely, the proportion of non-target contributor cells sorted into each fraction (6). In this study, the gates were set to minimize contributions from non-target cell populations in each fraction, with the expectation that the number of cells collected into the sorted fractions would be lower than the total number of cells and/or DNA present in the unsorted mixture sample). From these gate positions, we expected the female profile to be enriched in the right fraction, while the male profile was would be enriched in the left fractions. Analysis of unmixed donor cell populations indicated that the middle fraction would represent a mixture of both contributor cell populations.

Results showed that each sorted fraction contained alleles consistent with both male and female contributors. The number of sorted cells from each fraction and the resulting DNA yield are provided in Table S1. However, there were a greater number of alleles attributed to the male in the left and middle sorted fractions and conversely, a greater number of female alleles in the right fraction (Figure 5). It should be noted that female skin cell reference samples for the first two mixtures presented in this study showed extraneous alleles not attributable to the contributor (marked with an asterisk in Figures 4–7). The presence of extraneous alleles is not unexpected in a single contributor epithelial skin cell reference sample given the provenance of the sample and the low template status. Subsequent quantitative probabilistic modeling analyses were applied to the female profiles to capture the uncertainty of those genotypes when comparisons were performed between female donors and the pre-and post-sort fractions instead of just using each female donor’s single source genotype. This was not necessary for the male donors since the profiles were single source.

**Figure 4.**
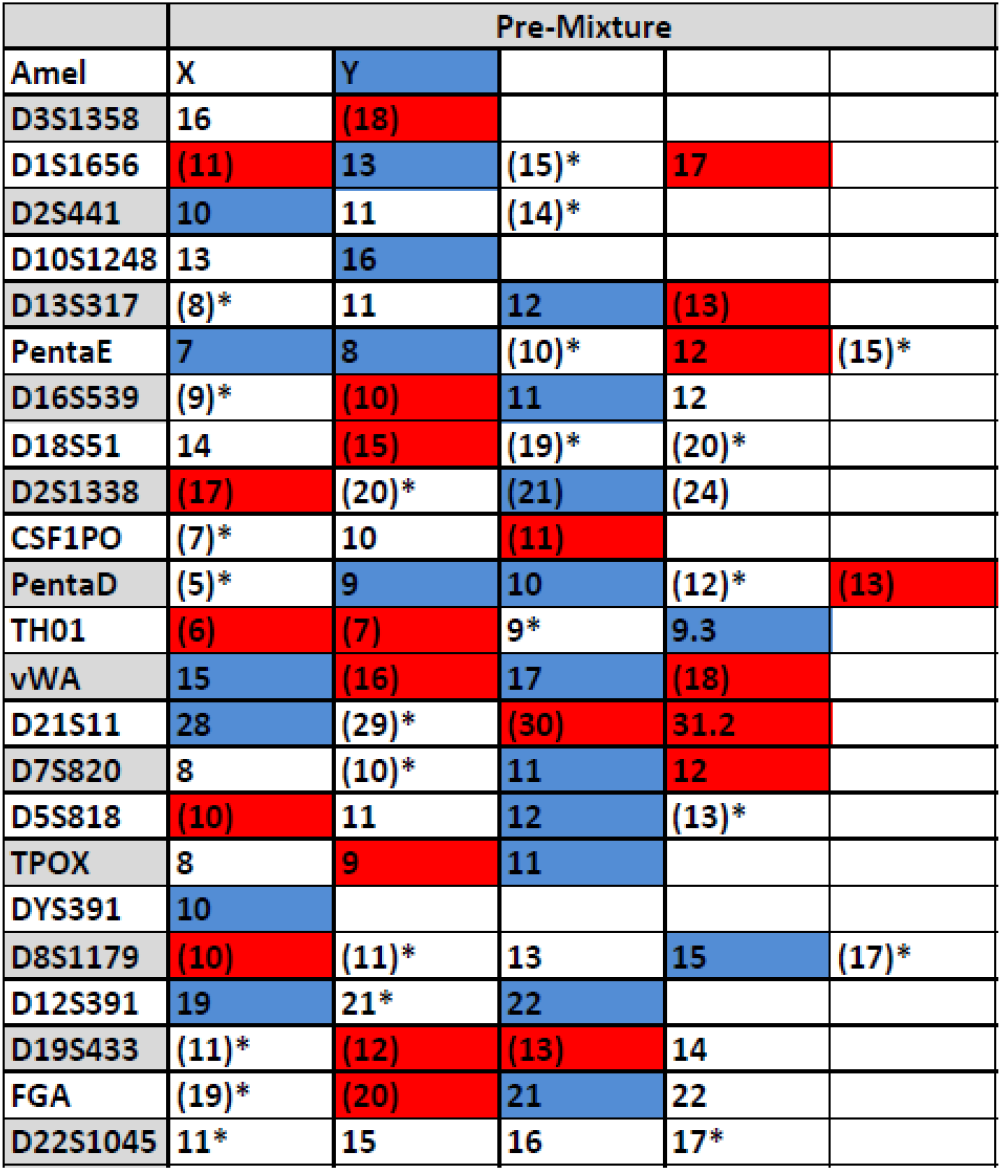
PowerPlex^®^ Fusion STR typing profile of the pre-sorted fraction of Mixture 1. Alleles belonging to the male contributor are highlighted in blue and alleles belonging to the female contributor are highlighted in red. Shared alleles that can be attributed to both the male and female contributor are uncolored. An asterisk (*) denotes an allele belonging to an additional contributor or pronounced stutter. Parentheses denote a minor, lower RFU allele.

**Figure 5.**
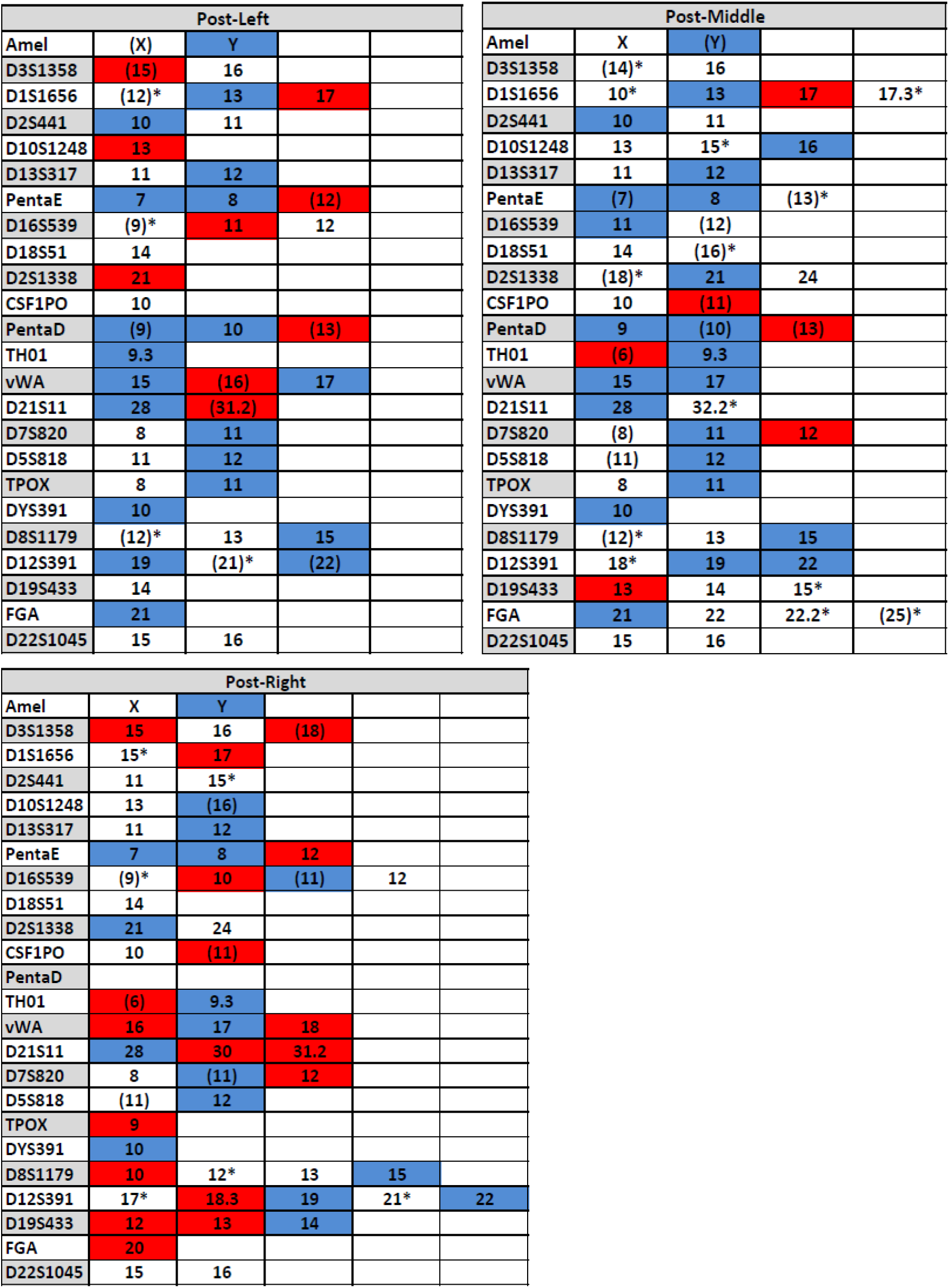
PowerPlex Fusion STR typing profile of the post-left (top left), post-middle (top right), and post-right (bottom) fraction of Mixture 1. Alleles belonging to the male contributor are highlighted in blue and alleles belonging to the female contributor are highlighted in red. Shared alleles that can be attributed to both the male and female contributor are uncolored. An asterisk (*) denotes an allele belonging to an additional contributor or pronounced stutter. Parentheses denote a minor, lower RFU allele.

This analysis suggests that male and female epithelial skin cells are enriched in sorted cell fractions compared to the original cell mixture. Next, we performed a quantitative assessment of cell population enrichment using TrueAllele^®^ Casework (TA) analysis. Results from the unsorted mixture showed statistical support for both the male and female contributors (Table 1). Following probe hybridization and cell sorting, the male donor profile had a log(LR) of 8.9791, while the female donor profile had a log(LR) of 1.0887 in the left sorted fraction. This supports inclusion of the male contributor in this fraction, whereas the female contributor cannot be conclusively associated (note that the inconclusive range is ±2.0 log(LR)) or it could be considered limited support if following the recommendations of the SWGDAM Ad Hoc Working Group on Genotyping Results Reported as Likelihood Ratios (20). The analysis of the sorted middle fraction showed high log(LR) value for the male contributor (12.2969 or 1.98 trillion time more likely) indicating strong statistical significance supporting the male contributor in this profile. On the other hand, a negative log(LR) value was produced for the female contributor indicating no statistical support for association with this fraction. Results from the right fraction showed evidence of both female and male contributor profiles (log(LR)s of 4.1185 and 3.7525, respectively). The log(LR) values for complete matches to the contributors are 25.6207 for the male and 25.8918 for the female contributor. However, note that complete matches would not be expected for such a challenging, low template, mixture sample types.

**Table 1.** TrueAllele ^®^ Casework analysis for Mixture 1, consisting of male and female epithelial skin cells. The logarithm of the likelihood ratio (log(LR)) values, regarding the statistical support of female and male STR profiles as contributors to the pre-sort mixture, post-sort middle fraction, and post-right fraction, are displayed. The log(LR) values of the male and female donor profiles as perfect matches to themselves are also displayed on the right side of the table. Log (LR) values less than 0 denote no statistical support.

| **log(LR) values** |  |  |  |  |  |
| --- | --- | --- | --- | --- | --- |
|  | **Pre-Sort** | **Post-Left** | **Post-Middle** | **Post-Right** | **Donor** |
| **Male** | 13.821 | 8.9791 | 12.2969 | 3.7525 | 25.6207 |
| **Female** | 3.1022 | 1.0887 | <0 | 4.1185 | 25.8918 |

### Fluorescence Activated Cell Sorting and DNA Profiling: Mixture 2

The gating criteria for the second mixture is shown in Figure 3 (right panel). Flow cytometry analysis of single donor cell solutions indicated that contributor ratio was ~1.1: 1 (M:F). DNA analysis of the unsorted mixture showed a mixture of male and female alleles (Figure 6). Although three different sorted cell fractions were initially collected (left, middle, and right), the cell solution sorted into the left fraction was compromised during sample handling and could not be analyzed. Qualitative analysis of STR profiles in the middle fraction showed more male contributor alleles, and at higher peak heights compared to alleles from the female contributor. Interestingly, a number of male alleles were detected in the after-sorting in the middle fraction, that were not observed in the STR profile of the unsorted mixture. While the unsorted mixture displayed many alleles consistent with the male contributor, the accumulation of missing allelic information led to the lack of a sufficient statistical support for the male contributor to the unsorted mixture (one locus drop-out, three allelic drop-outs and 8 allelic dropouts with the sister allele shared with the female contributor [data not shown, Figure 6 and Table 2]). This suggests selective enrichment of the male contributor. Given the low template of the epithelial skin cell samples and many peaks within the stochastic range, it is not unexpected that pre-and post-sort sample profiles may display allelic and locus drop-out. Conversely, the right fraction showed more female contributor alleles compared to the male contributor (Figure 7) in the unsorted mixture and the sorted middle fraction.

**Figure 6.**
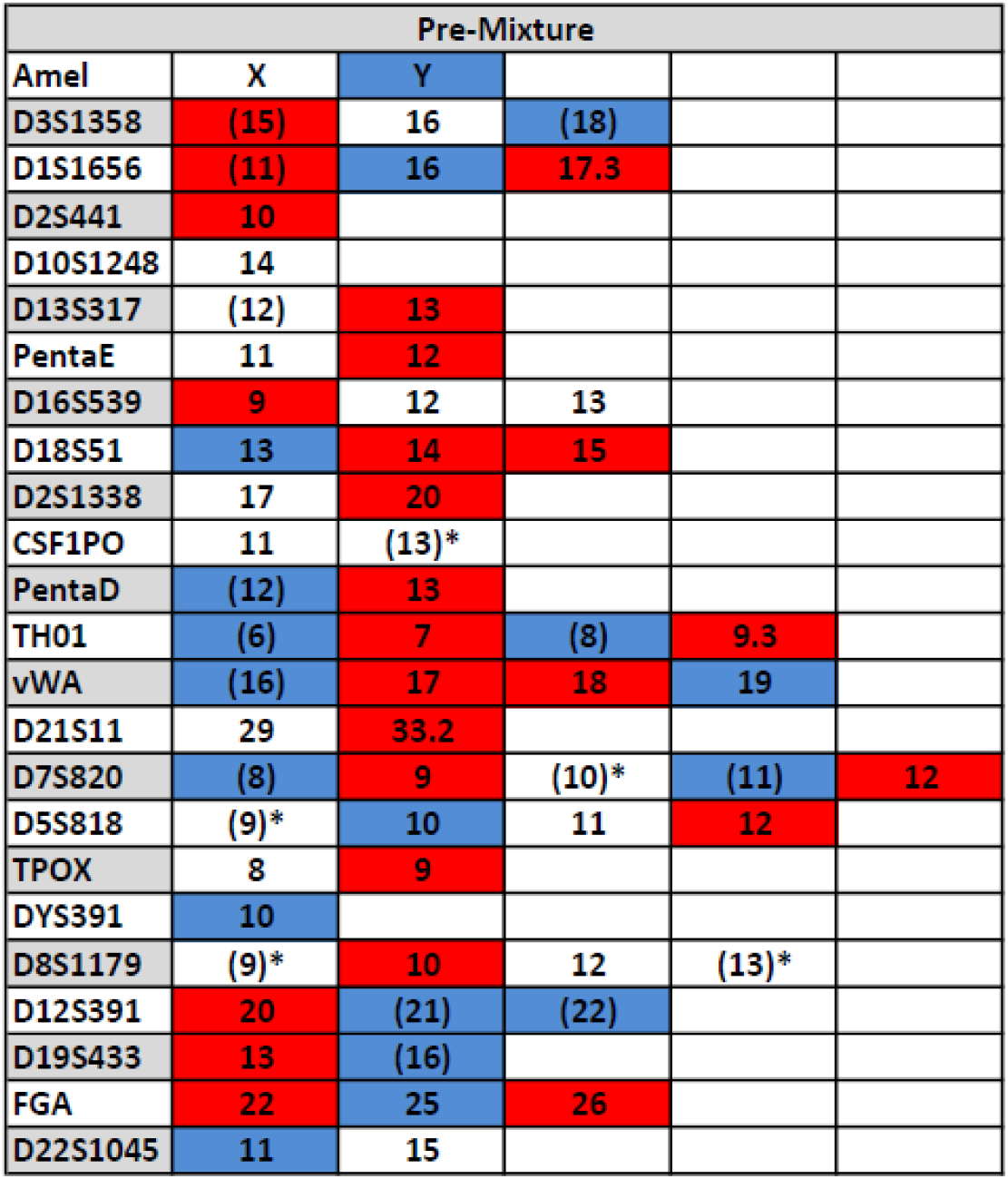
PowerPlex^®^ Fusion STR typing profile of the pre-sorted fraction of Mixture 2. Alleles belonging to the male profile are highlighted in blue and alleles belonging to the female contributor are highlighted in red. Shared alleles that can be attributed to both the male and female contributor are uncolored. An asterisk (*) denotes an allele belonging to an additional contributor or pronounced stutter. Parentheses denote a minor allele.

**Figure 7.**
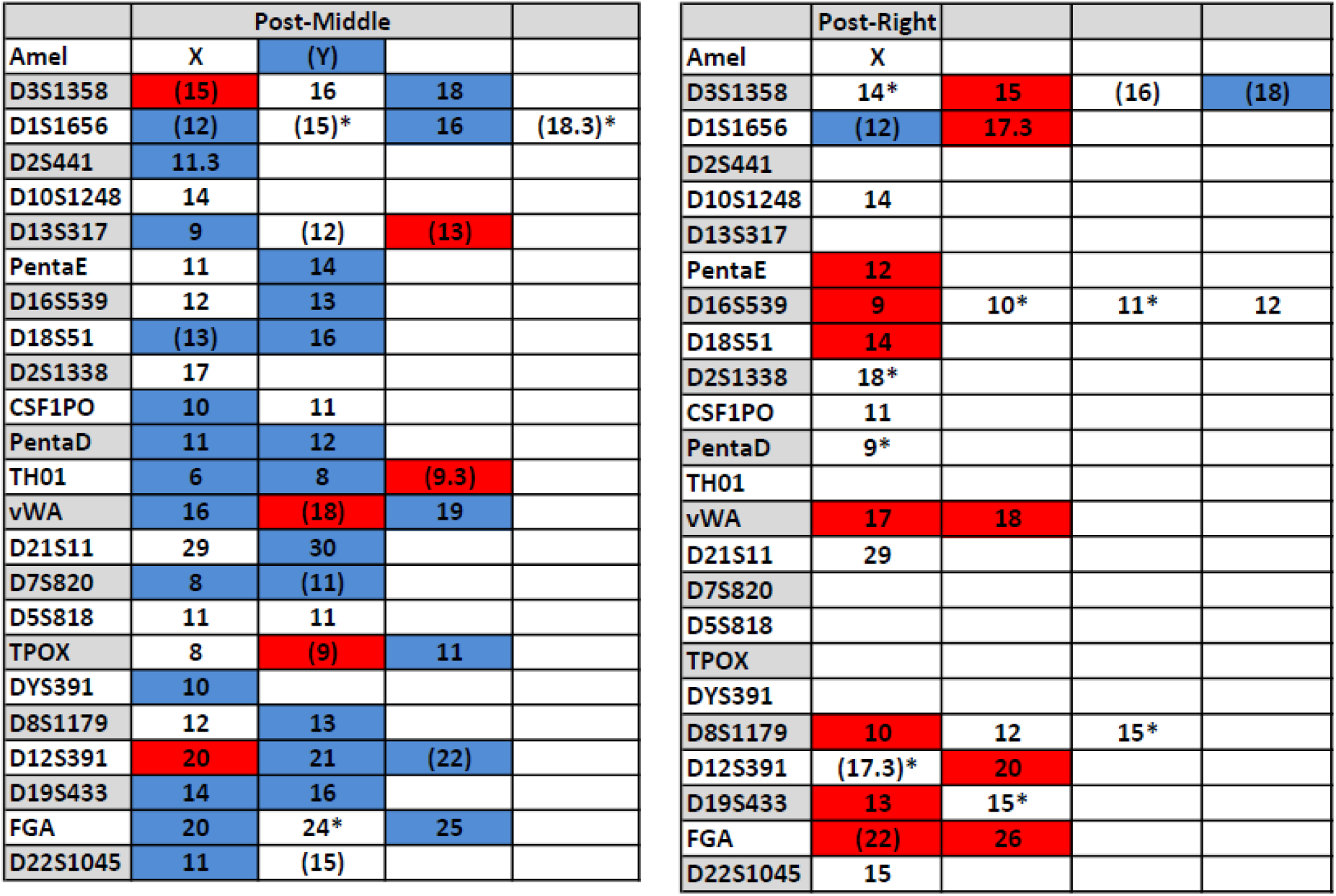
PowerPlex Fusion STR typing profile of the post-middle and post-right fraction of Mixture 2. Alleles belonging to the male profile are highlighted in blue and alleles belonging to the female contributor are highlighted in red. Shared alleles that can be attributed to both the male and female contributor are uncolored. An asterisk (*) denotes an allele belonging to an additional contributor or pronounced stutter. Parentheses denote a minor allele.

**Table 2.** TrueAllele ^®^ Casework analysis for Mixture 2, consisting of male and female epithelial skin cells. Log Likelihood Ratio (log(LR)) values, regarding the statistical support of female and male STR profiles as contributors to the pre-sort mixture, post-sort middle fraction, and postright fraction, are displayed. The log (LR) values of the male and female donor profiles as perfect matches to themselves are also displayed on the right side of the table. Log(LR) values less than 0 denote no statistical support.

|  | **log(LR) values** |  |  |  |
| --- | --- | --- | --- | --- |
|  | **Pre-Sort** | **Post-Middle** | **Post-Right** | **Donor** |
| **Male** | <0 | 19.5523 | <0 | 26.9279 |
| **Female** | 14.5475 | <0 | 2.9306 | 25.6499 |

Quantitative analysis with TA also indicated enrichment of male and female profiles in the middle and right fractions, respectively. The log(LR) for the male contributor in the middle fraction was 19.5523 (i.e., 35 quintillion times, more likely for the male profile in this fraction compared to coincidence) (Table 2). Analysis of the unsorted mixture indicated no statistical support for the male contributor in the unsorted mixture (log(LR) <0), consistent with the minimal number of unique male alleles observed and excessive allelic and locus drop-out. The drastic increase in log(LR) value after sorting is strong evidence for selective enrichment of male epithelial cells in this fraction.

The right fraction had a log(LR) of 2.9306 for the female contributor, which indicates that there is some statistical support for the female to this sorted fraction. As expected, there was high statistical support for the female as a contributor to the pre-sort mixture, but the post-right fraction was at a low level, with significant allelic drop-out of alleles attributable to the female. The female donor showed a reproducible negative log(LR) for its association to the post-middle fraction. The log(LR) values for perfect matches to the contributors are 26.9279 for the male contributor and 25.6499 for the female contributor. Some of the large discrepancy in the log(LR) values between the pre-sort mixture and the post-right fraction may be due the presence of eDNA in the pre-sort fraction.

### Fluorescence Activated Cell Sorting and DNA Profiling: Mixtures 3, 4

As a preliminary test for this cell separation workflow on unknown mixture samples, we created two additional mixtures composed of one male and one female donor. However, unlike the previous two mixtures, single source cell populations from each donor were not analyzed prior to sorting in order to guide placement of the sorting gate. Fluorescence histograms for both mixtures show similar distributions and median fluorescence values (2-3×10^3^ RFU; Figures S4, S5). A single sorting gate was used to collect two fractions (‘post-left’ and ‘post-right’) (Table S1). Quantitative analysis of the unsorted mixture samples showed strong support for both the male and female profiles (Tables S2, S3), however, the male to female ratio was estimated by probabilistic modeling to be ~9:1. For Mixture 3, there was evidence that the male profile was highly enriched in both the right and left fractions, log(LR) 24.537 and 29.0711, respectively (Table S2). However, there was little support for the female profile in either fraction, log(LR) <0 and 0.6731 in the right and left fraction respectively. For Mixture 4, there was strong statistical support for the male and female profiles in both the right and left sorted cell fractions (Table S3). Differences were observed in the magnitude of the log (LR) values between the sorted cell fractions with the male displaying a higher log(LR) association than the female in the right fraction (13.374 vs. 7.8387) and the female higher than the male in the left fraction (12.263 vs. 6.9911), Table S2. The higher and lower log(LR) associations correlate to the estimated mixture weight proportions of the contributors in the two fractions (data not shown).

## Discussion

The goal of this effort was to develop a method for differentiating contributor cell populations based on interactions with hormone-specific antibody probes and then investigate whether differences could be exploited to separate each mixture component prior to DNA profiling. Overall, antibody probes targeting testosterone and dihydrotestosterone molecules showed strong interactions with trace cell populations from all individuals sampled in this study. Variation in antibody interactions were also observed between contributor cell populations and did not strictly correlate with sex of the individual. In fact, for the set of donor cell samples used in this study, epithelial cell populations from female contributors often showed higher affinity for testosterone and dihydrotestosterone antibody probes in comparison to epithelial cell populations from male contributors. There are several possible factors contributing to this. First, hormone levels in serum can vary between individuals of the same sex due to age, genetics, or environmental factors (11–13,21). Additionally, the abundance of hormone targets like testosterone or dihydrotestosterone in terminally differentiated epidermal cells is largely unknown but is likely to differ from well documented levels observed in serum and gonadal tissue across males and females. Within epidermal tissue, hormone molecules may be influenced by contributor-specific biological factors such as cell turnover rate, epidermal thickness, and the biochemical profile of the epidermis (22–24). Probes may also cross-react with other steroid or steroid-like molecules within epidermal cells (25), however, for the antibodies utilized in this study, the reported cross-reactivity with other non-testosterone analog cholesterol-based molecules (e.g. estradiol) was less than 1% and testing of the anti-testosterone antibody with purified testosterone and estradiol confirmed an undetectable or extremely low level of crossreactivity (data not shown). Sampling factors for this study may have also lead to differences in cell populations, since a few donors collected cells from only one part of face/neck area (ears) whereas others collected cells from only the outer skin of the nose (See Methods) to minimize possible interferences from skin/makeup substances. Further optimization of the antibody binding conditions may be necessary in order to enhance the detected differences in male and female epithelial cell populations. This could include careful optimization of antibody binding concentrations, focusing on specific sub-populations of epidermal cells within the touch sample or investigating other orthogonal hormone targets in combination with testosterone and/or dihydrotestosterone (e.g., estradiol).

Results from sorting contributor cell populations based on differences in antibody labelling showed some selective enhancement of male and female contributor profiles. In the first two mixtures where male and female cell components were present in approximately equivalent ratios (Mixtures shown in Figure 3), TA derived contributor profiles could be resolved from separated cell populations with statistical significance. For other mixtures tested in a ‘blind’ fashion whereby donor cell populations were not analyzed initially to guide gate selection, the male component was resolved in both the left and right sorted cell fractions whereas the female component was not detected in either fraction after sorting (Mixture 3; Figure S4). For Mixture 4, both contributor profiles were resolved in each sorted cell fractions with weaker evidence for enrichment of female and male profiles in the left and right fractions respectively (Mixture 4; Figure S5).

The limited enrichment of contributor profiles in the latter two ‘blinded’ mixture samples suggests that the sorting gates may need further optimization before this approach can be applied to unknown mixture samples. This is not unexpected given the differences in antibody binding efficiency observed between contributors (Figure 2). Consequently, the fluorescence regions of the sorting histogram that are most likely to selectively isolate one contributor are likely to vary depending on the donor cell populations present. Possible strategies to increase the efficacy of unknown sample sorting include isolating more than two cell fractions (as demonstrated in the first two mixtures, Figure 3), or reducing the potential for clumping/aggregation of cells from different contributors during sorting by adding surfactants or changing the buffer chemistry of the initial cell mixture. Additionally, further optimization of antibody binding conditions could contribute to a more reproducible and definitive separation of the sexes into post-sort fractions. Another strategy is to shift the position of the sorting gate to capture a narrower subset of the original cell population, thereby reducing the total number of cells collected in a given fraction and increasing the potential selectivity for a specific contributor.

However, one of the primary obstacles for front end separation techniques on trace biological samples may be the proportion of extracellular versus intracellular DNA. For cell populations derived from sloughed epidermal cells, the quantity of DNA that is physically associated with cells and may be especially limited (6) with the majority of total DNA comprised of extracellular or cell-free DNA (eDNA/cfDNA) as reported in previous studies (26–28). During the cell separation process, this DNA would be dissociated from the cells and may be partitioned with the effluent buffer and not collected with either sorted cell fraction. Although the relative proportion of eDNA was not quantified in these mixture samples, this would likely explain the low DNA yields in each sorted fraction given the respective cell counts and when compared to the pre-sort fractions (Table S1). Future efforts should focus on integrating information from extracellular DNA fractions of trace mixtures with contributor profiles obtained from FACS sorted cells. This could be accomplished by either capturing free DNA during the cell separation process (e.g., in-line filter) or collecting eDNA as part of a pretreatment procedure prior to antibody labelling and cell sorting.

## Conclusion

In summary, the results from this study show that trace epithelial cell populations can be labeled through the use of antibody probes targeting testosterone and dihydrotestosterone sex hormones within cell populations and that differences in antibody binding efficiency between male and female contributors can potentially be used to differentiate and ultimately separate cell populations. Coupling hormone-specific antibody probes with fluorescence activated cell sorting of two-person mixtures showed successful enrichment for contributor cell populations for two different, two-person mixtures. When applied to two additional mixtures in a blind fashion, evidence for successful enrichment was observed in one of the two, albeit not as pronounced as was observed in the first two mixtures presented. The other of the two additional mixtures showed a large contributor proportion disparity, 1:9, in the pre-sort fraction and did not result in enriching male and female contributors into distinct fractions. This indicates that sorting efficiency at this point in the development of this procedure may depend partly on the nature of the donor cell populations present. Overall, although future research can improve the efficiency of separation and subsequent resolution of contributor DNA profiles, our results indicate that hormone-specific antibodies may be a useful tool to label epidermal cell populations, one of the most challenging cell types associated with trace biological evidence. In addition to the potential use for front-end cell separation, differences in binding efficiency of hormone-specific antibody probes can potentially be used to presumptively detect the presence of multiple contributor cell populations.

## Acknowledgements

The authors gratefully acknowledge Julie Farnsworth for providing technical assistance for this project.

