## Supplemental Figures and Tables for "Testing Hormone-specific Antibody Probes for Presumptive Detection and Separation of Contributor Cell Populations in Trace DNA Mixtures"

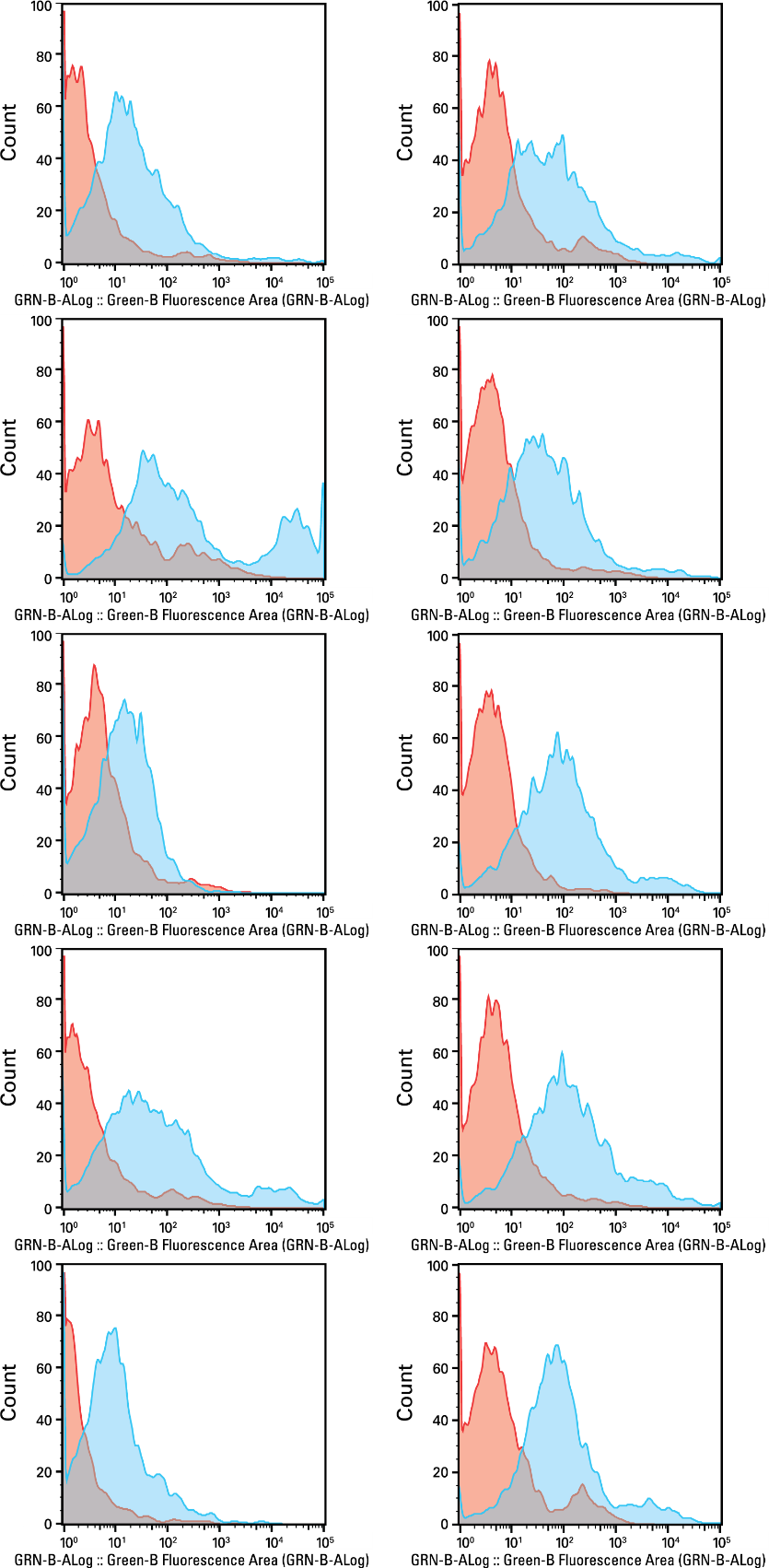

**Figure S1**. Fluorescence histograms of male epithelial skin cell populations hybridized to anti-testosterone and anti-dihydrotestosterone antibodies (blue) and unstained (red) from the same contributor. Results from 10 different male contributors are shown. The X-axis shows fluorescence intensity, while the Y-axis shows the number of events analyzed for each sample.

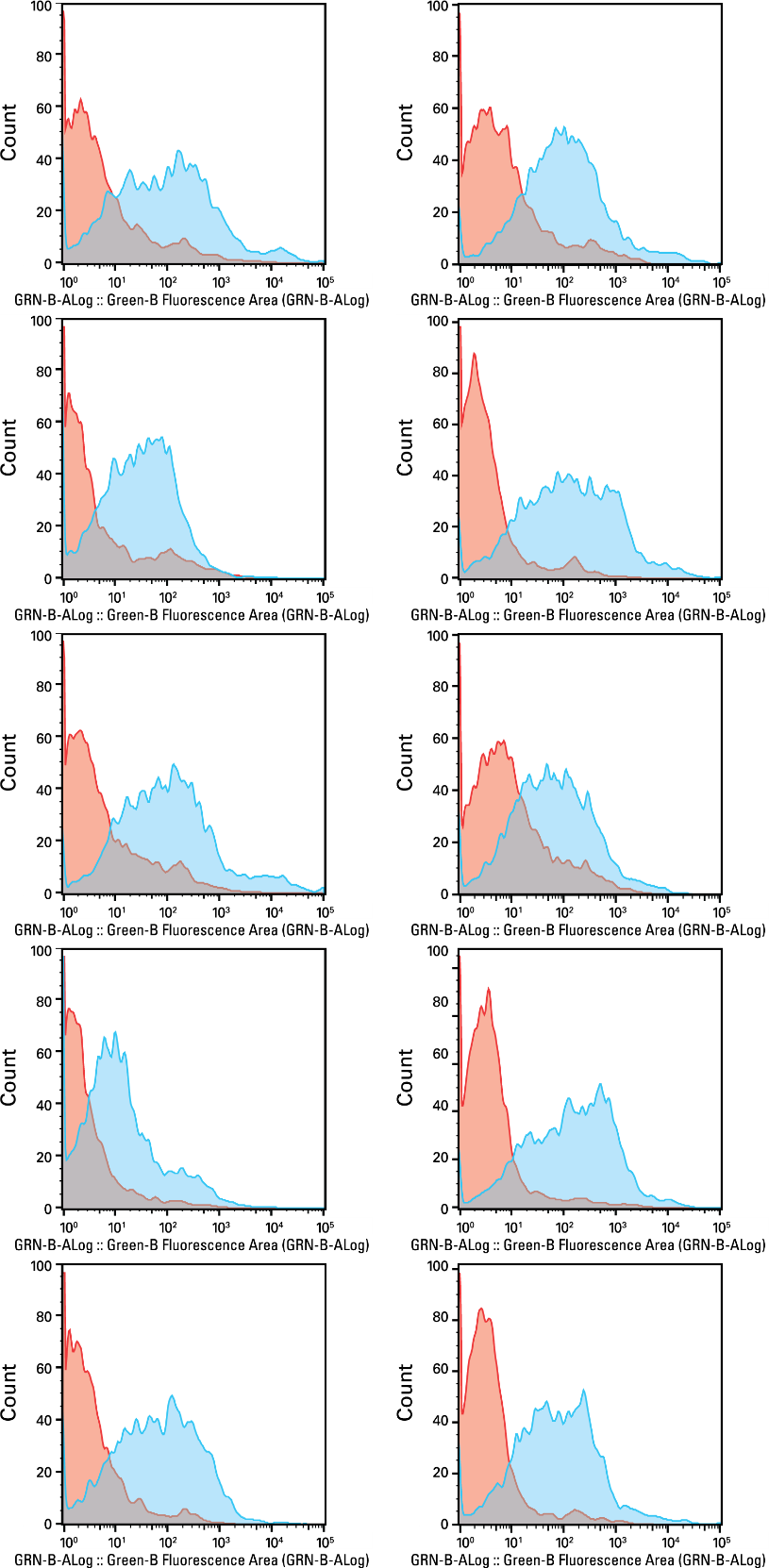

**Figure S2.** Fluorescence histograms of female epithelial skin cell populations hybridized to anti-testosterone and anti-dihydrotestosterone antibodies (blue) and unstained (red) from the same contributor. Results from 10 different female contributors are shown. The X-axis shows fluorescence intensity, while the Y-axis shows the number of events analyzed for each sample.

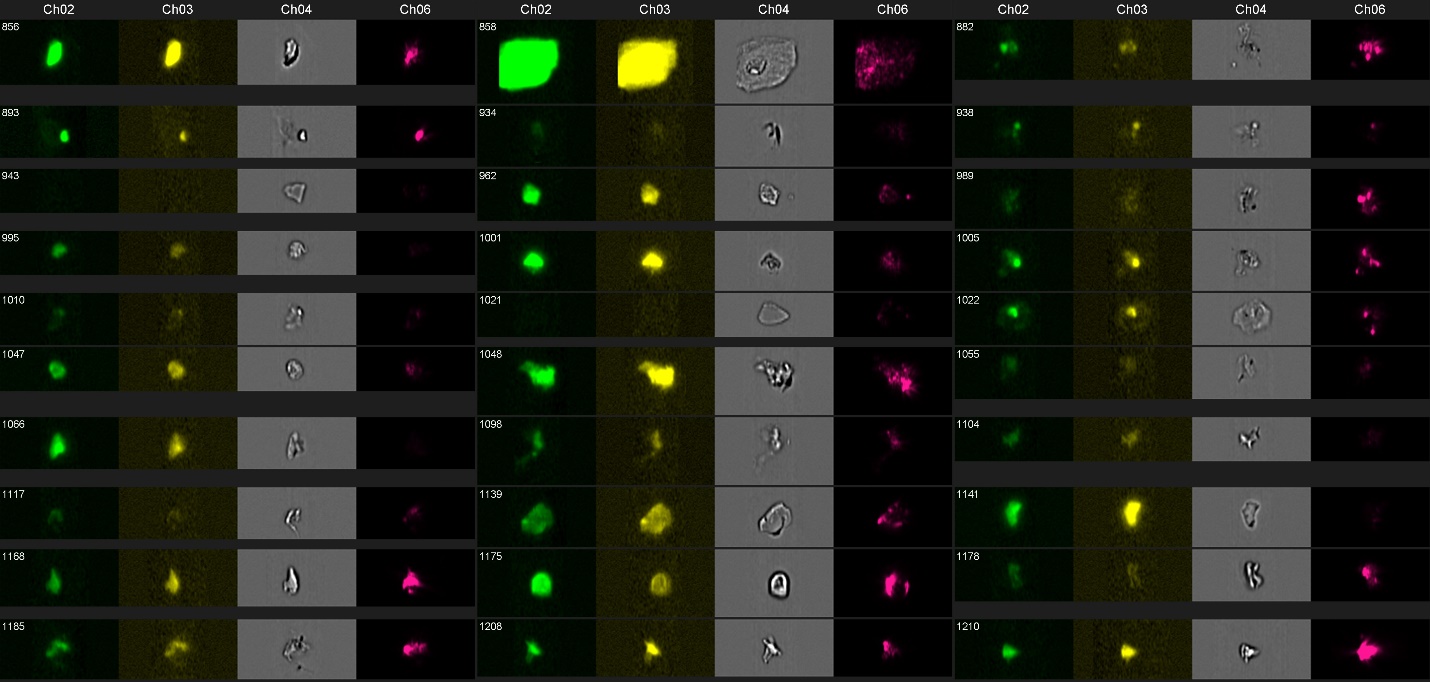

**Figure S3**. Image galleries of events analyzed from male epithelial cell populations after labeling with testosterone-specific antibody probe. Full image galleries from both male and female contributor cell populations are provided in associated data repositories

(<https://figshare.com/s/999dc144d7f68b13a699>)

**Table S1. Sorted Cell Populations and DNA Yield for Mixtures**

*Mixture 1*

Left fraction	Middle fraction	Right fraction

\# of sorted cells/debris	4,205	2,179	449

DNA Yield	36 pg	100 pg	48 pg

*Mixture 2*

Left fraction	Middle fraction	Right fraction

\# of sorted cells/debris	4,144	1,939	981

DNA Yield	(fraction missing)	88 pg	120 pg

*Mixture 3 (no quantification – straight to amp)*

Left fraction	Right fraction

\# of sorted cells	4,096	765

DNA Yield

*Mixture 4*

Left fraction	Right fraction

\# of sorted cells	4,062	829

DNA Yield

|  | **log(LR) values** | | | |
| --- | --- | --- | --- | --- |
|  | **Pre-Sort** | **Post-Left** | **Post-Right** | **Donor** |
| **Male** | 29.015 | 29.071 | 24.537 | XX |
| **Female** | 16.819 | 0.673 | <0 | XX |

**Table S2**. TrueAllele ^®^ Casework analysis for Mixture 3, consisting of male and female epithelial skin cells. The logarithm of the likelihood ratio (log(LR)) values, regarding the statistical support of female and male STR profiles as contributors to the pre-sort mixture, post-sort left fraction, and post-right fraction, are displayed.

|  | **log(LR) values** | | | |
| --- | --- | --- | --- | --- |
|  | **Pre-Sort** | **Post-Left** | **Post-Right** | **Donor** |
| **Male** | 29.015 | 29.071 | 24.537 | XX |
| **Female** | 16.819 | 0.673 | <0 | XX |

|  | **log(LR) values** | | | |
| --- | --- | --- | --- | --- |
|  | **Pre-Sort** | **Post-Left** | **Post-Right** | **Donor** |
| **Male** | 29.015 | 29.069 | 24.482 | 29.480 |
| **Female** | 14.354 | -0.668 | -0.948 | 29.083 |

|  | **log(LR) values** | | | |
| --- | --- | --- | --- | --- |
|  | **Pre-Sort** | **Post-Left** | **Post-Right** | **Donor** |
| **Male** | 18.731 | 6.847 | 12.848 | 29.159 |
| **Female** | 15.480 | 11.611 | 7.758 | 35.075 |

**Table S3.** TrueAllele ^®^ Casework analysis for Mixture 4, consisting of male and female epithelial skin cells. The logarithm of the likelihood ratio (log(LR)) values, regarding the statistical support of female and male STR profiles as contributors to the pre-sort mixture, post-sort left fraction, and post-right fraction, are displayed.

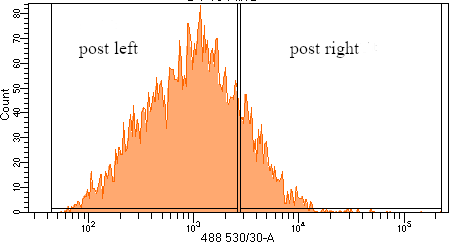

**Figure S4**. Cell population histogram and sorting gates for Mixture 3. Sorting gate used for separating right and left fraction shown (~2600 RFU).

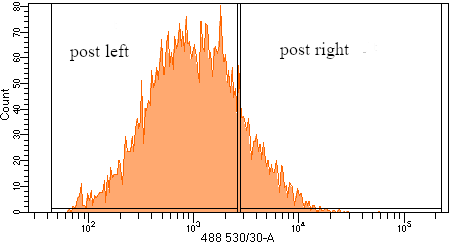

**Figure S5**. Cell population histogram and sorting gate for Mixture 4. Sorting gate used for separating right and left fraction shown (~2600 RFU).
